# PAX2 Induces Tubular-Like Structures in Normal and Ovarian Cancer Cells

**DOI:** 10.1101/2020.01.14.906438

**Authors:** Kholoud Alwosaibai, Ensaf Munawer Al-Hujaily, Salmah Alamri, Kenneth Garson, Barbara C. Vanderhyden

## Abstract

In adult tissues, PAX2 protein is expressed in normal oviductal epithelial cells but not in normal ovarian surface epithelial cells. Studies have reported that PAX2 is expressed in a subset of serous ovarian carcinoma cases but the role of PAX2 in the initiation and progression of ovarian cancer remains unknown. The aim of this study was to understand the biological consequences of *Pax2* expression in normal and cancerous mouse epithelial (MOSE) cells. We found that *Pax2* overexpression in both normal and cancerous ovarian epithelial cells induced the formation of vascular channels both *in vitro* and *in vivo*. The results indicate a possible contribution of PAX2 to ovarian cancer progression by increasing the vascular channels to supply nutrients to the tumor cells.

## Introduction

Angiogenesis is required to establish the blood supply needed for tumor growth. Anti-angiogenesis treatment approaches have been used widely to inhibit the neovascularization, but the results have been unsatisfactory in invasive and aggressive cancers (1,2). The growth of tumors even in the presence of anti-angiogenesis drugs indicates that tumor cells may have other ways to be supplied with blood. Antiangiogenesis therapies, such as anti-VEGF antibodies, target only endothelial cell proliferation, whereas cancerous cells may proliferate and form *de novo* vascular channels connected to the endothelial-lined vasculature.

In 1999, Maniotis et al. observed aggressive melanoma cells forming a blood vessel-like pattern in 3D culture and, when the cells were injected into mice they formed tumors that contained vessel-like structures. This formation of vessel-like structures was termed vasculogenic mimicry (VM) (3). Interestingly, red blood cells were detected inside the vessels while the cells lining these vessels were negative for the endothelial markers CD31 and CD34 (4). The tumor cells were organized in a pattern that was supported by remodeling extracellular matrix, as determined by Periodic Acid-Schiff (PAS) staining to label the carbohydrates in the remodeling extracellular matrix (5).

In more recent studies, CD31-/PAS+ vessels formed by tumor cells have been reported in several studies to demonstrate VM *in vitro* and *in vivo* (6–8) and some attempts have been made to determine the molecular mechanisms controlling VM formation (9). In aggressive breast cancer cells, cyclooxygenase-2 (COX2) can induce VM formation in three-dimensional culture, and knockdown of COX2 decreased VM formation (10). The ovarian cancer cell line SKOV3 has been reported to form VM *in vitro* and targeting CD147 in these cells caused a significant down-regulation in VM formation (11). CD147 protein is an inducer of extracellular matrix metalloproteinase (MMP) (12). Overexpression of MMP has been linked previously with VM incidence in ovarian cancer cell lines SKOV3 and OVCAR3, where MMP expression contributed to the extracellular matrix remodeling (13).

Ovarian cancer cell lines have been reported to form VM both in matrigel and after being xenografted in mice (13,14). The vascular channels formed by SKOV3 and OVCAR3 cells were matrix-enriched and endothelial cell-independent, and the cells lining the vascular channels were positive for cytokeratin (13). Interestingly, blood cells were detected inside the vascular channels formed *in vivo* (3,5). To identify the enriched extracellular matrix (ECM), tumors formed by the ovarian cancer cells were stained for laminin, which was found to be abundantly expressed by the cancer cells and present in the remodeling ECM. How the ovarian cancer cells promote VM is unknown, but some ovarian cell lines express endothelial markers *de novo* and it has been suggested that cancer cells may differentiate into endothelial cells to promote angiogenesis (15). The SKOV3 cell line has the ability to form VM in 3D culture, which may be achieved by its weak expression of the endothelial marker CD31 that is evident on the cell surface in monolayer cultures and in the VM formed by the cells (16).

Ovarian carcinomas are heterogeneous with few genetic mutations but considerable genomic aberrations associated with altered gene expression. PAX2 has been reported to be up-regulated in some types of ovarian cancer, including high-grade serous carcinoma (17), but its role in promoting tumor progression is not clear. PAX2 plays very important roles during embryogenesis, driving epithelial differentiation during urogenital tract development (18). PAX2 is also thought to induce cell elongation and tubule formation during embryonic development in different types of organs such as inner ear, neural tubes and genital tract (19). We have previously demonstrated that PAX2 is critical for maintaining epithelial cell differentiation in adult reproductive tissues (20). In the kidney, PAX2 plays a very important role in the development of the collecting duct, where it induces branching morphogenesis (21,22). This was confirmed in PAX2 mutant mice, which have decreased ability to form branched collecting duct compared to PAX2 wild type mice (22). The ability of PAX2 to promote the formation of branching structures by epithelial cells led us to hypothesize that PAX2 expression in ovarian cancers may promote VM to enhance tumor growth.

## Methods

### Cell lines

Normal mouse ovarian surface epithelial (MOSE) cells (M1102) were isolated from the ovarian surface of 6-week-old FVB/N mice and cultured in DMEM media supplemented with 10% fetal bovine serum (FBS) as described previously (23). Ovarian cancer cells (RM) were derived from immortalized MOSE cells that were transduced with c-*Myc* and *K-Ras* (KRAS^G12D^) and maintained in DMEM media supplemented with 10% FBS (24). Ovarian cancer cells SKOV3 were maintained in DMEM media supplemented with 10% FBS. Human Umbilical Vein Endothelial Cells (HUVEC) were obtained as a gift from Dr. Christina Addison. HUVEC cells were maintained in EGM-2 media (LONZA, Switzerland) supplemented with growth factors and 10% FBS.

### Animals

Animal experiments were conducted in accordance with the guidelines of the Canadian Council on Animal Care under a protocol approved by the University of Ottawa’s Animal Care Committee. Eight-week old SCID mice were injected into the peritoneal cavity with ovarian cancer cells. Three mice were euthanized when they reached humane endpoint and the tumor tissues were collected for immunohistochemistry (25).

### Lentiviral vectors

The murine *Pax2* cDNA, corresponding to the pax2-b variant, was cloned from the murine oviduct and inserted into pWPI (Addgene plasmid #12254) to generate the lentivirus expression vector (WPI-Pax2-IRES-eGFP) as described previously (25). Lentiviral vectors were prepared by co-transfection of vector plasmids with packaging plasmid pCMVR8.74 (Addgene plasmid #22036) and the ecotropic envelope expression plasmid, pCAG-Eco (Addgene plasmid 35617) into 293T cells as described previously (26). Plasmids pWPI and pCMVR8.74 were gifts from Didier Trono and plasmid pCAG-Eco was a gift from Arthur Nienhuis and Patrick Salmon.

### PAX2 overexpression and knockout

M1102 and RM cells were infected with either the control lentiviral vector, WPI, or WPI-Pax2-IRES-eGFP to generate cell lines M1102-WPI, M1102-PAX2, RM-WPI and RM-PAX2, as described in (25). As an additional control, the *Pax2* expression cassette was deleted from M1102-PAX2 cells following Cre-mediated recombination between *loxP* sites resident in both the 5’ and 3’ LTRs of the integrated WPI-Pax2-IRES-eGFP provirus. Cre recombinase was delivered by the transient infection of M1102-PAX2 cells with adenovirus expressing Cre recombinase (AdCre; Vector Development Laboratory, Houston, Texas) to form M1102-PAX2-AdCre cells (25).

### Culture in matrigel

Chamber slides (8 wells; Thermo Fisher) were covered with 60 μl of cold growth factor-reduced matrigel (Trevigen, Gaithersburg, MD) and incubated at 37C° for 30 minutes to polymerize the matrigel. M1102 cells, SKOV3 and HUVEC were mixed with assay media (MEGM™ Mammary Epithelial Cell Growth Medium supplemented with 5% fetal bovine serum, 1 ng/ml EGF and 0.5 ng/ml of insulin, hydrocortisone, cholera toxin and gentamicin (all from lonza, Switzerland) and plated on top of the matrigel as described previously (27). The cells were covered with assay media containing 4% matrigel and incubated at 37°C for 48 hours. RM cells were similarly plated, however serum-free DMEM medium was used in place of the MEGM medium described. The formation of vascular channels was quantified by counting all tubule branches in each well.

### Immunofluorescence

Immunofluorescence was performed on cells grown in matrigel. The cells were fixed with 4% paraformaldehyde (Sigma-Aldrich, Missouri) for 20 minutes and permeabilized with 1:1 methanol and acetone for 15 minutes in 4C°. After blocking in 5% goat serum, cells were probed with anti-rabbit CD31 (1:100; abcam). Alexa Fluor goat anti-rabbit IgG (1:500, Carlsbad, CA) was used as secondary antibody. The cells in the matrigel slides were mounted with cover slips using Vectashield hard set mounting medium with DAPI (Vector Laboratories, Burlingame, CA) and the immunofluorescence images were visualized using an inverted fluorescence microscope (Axioskop 2 MOT plus, Zeiss) and Axiovision software.

### Immunohistochemistry (IHC)

Tumors were excised from euthanized mice and fixed in formalin overnight, then transferred to 70% ethanol. The tissue was paraffin-embedded, cut in 4 μm sections, deparaffinized and rehydrated in graded ethanol. For antigen retrieval, the tissues were heated in a microwave in antigen unmasking solution (Vector Laboratories). Tissue sections were blocked with Dako protein block, serum-free (Dako, Glostrup, Denmark) for 1 hour and then incubated overnight at 4C°with anti-rabbit CD31 antibody (1:100, Abcam) or 1 hour at room temperature for anti-rabbit PAX2 (1:1000, Santa Cruz). After washing in phosphate-buffered saline (PBS), the sections were incubated with ImmPRESS anti-rat secondary antibody (Vector Laboratories) or with Dako Envision system-HRP labeled polymer anti-rabbit antibodies (Dako; Burlington, ON) for 30 minutes. The immunoreactivity was imaged using the Aperio ScanScope system and analyzed using ImageScope software. Protein expression was quantified as positive pixels using Aperio Positive Pixel Count Algorithm software (Leica Biosystem, IL).

### Periodic Acid-Schiff (PAS) staining

PAS staining was used for either paraffin-fixed tissues or for 3D culture to stain the tubular structures. For paraffin-fixed tissues, the fixed tissues were cut, de-paraffinized, rehydrated and then heated for antigen retrieval. For vascular channels formed by MOSE and RM cells in 3D culture, the 8 well chamber slides were washed carefully with PBS, fixed with 2% paraformaldehyde for 15 minutes, and then washed with PBS three times. PAS staining was done using the Periodic Acid-Schiff kit (Sigma-Aldrich, Missouri) following the manufacturer’s protocol. Periodic acid (250 μl/well) was added to the cells, incubated at room temperature for 5 minutes, and then washed with PBS 3-5 times for 5 minutes each wash. Schiff stain (250 μl/well) was then added, the cells were incubated at room temperature for 15 minutes and then washed 3 times for 5 minutes each wash. All washing steps were with 500 μl/well of PBS using a pipet to drip the solution carefully on top of the matrigel.

### Statistical analyses

All experiments were performed a minimum of three times. Image Analyses were done using Aperio Positive Pixel Count Algorithm software. Statistical analyses were performed using GraphPad Prism software (GraphPad, La Jolla, CA). A Student t-test (two groups) or an ANOVA with Tukey’s post-test (multiple groups) was used to determine statistical significance (P< 0.05). Error bars represent the standard error of the mean.

## Results

### PAX2 induces vascular channels *in vitro* in MOSE cells

The overexpression and knockout of PAX2 in M1102 cells were confirmed using western blot assay (Fig 1A). To explore the possibility that PAX2 could induce vascular channel formation by normal epithelial cells, M1102-PAX2 cells were plated in 3D culture of matrigel, using HUVEC cells as a positive control. M1102–PAX2 cells were highly branched and elongated to form tubular channels that mimic the vascular channels formed by the endothelial cells (Fig 1). Interestingly, we found that the formation of tubular channels is highly dependent on cell density. The higher number of cells plated formed very organized branched vascular channels within 48 hours (Fig 1B and D). In contrast, the parental cells and vector control M1102 cells did not form these tubular channels and deletion of *Pax2* by adding AdCre to the cells decreased their ability to form tubular channels (Fig 1B). The statistical analysis of the tubule formation shows a significant increase in the number of tubular branches formed by M1102-PAX2 compared to the control groups (Fig1C).

**Fig 1:**
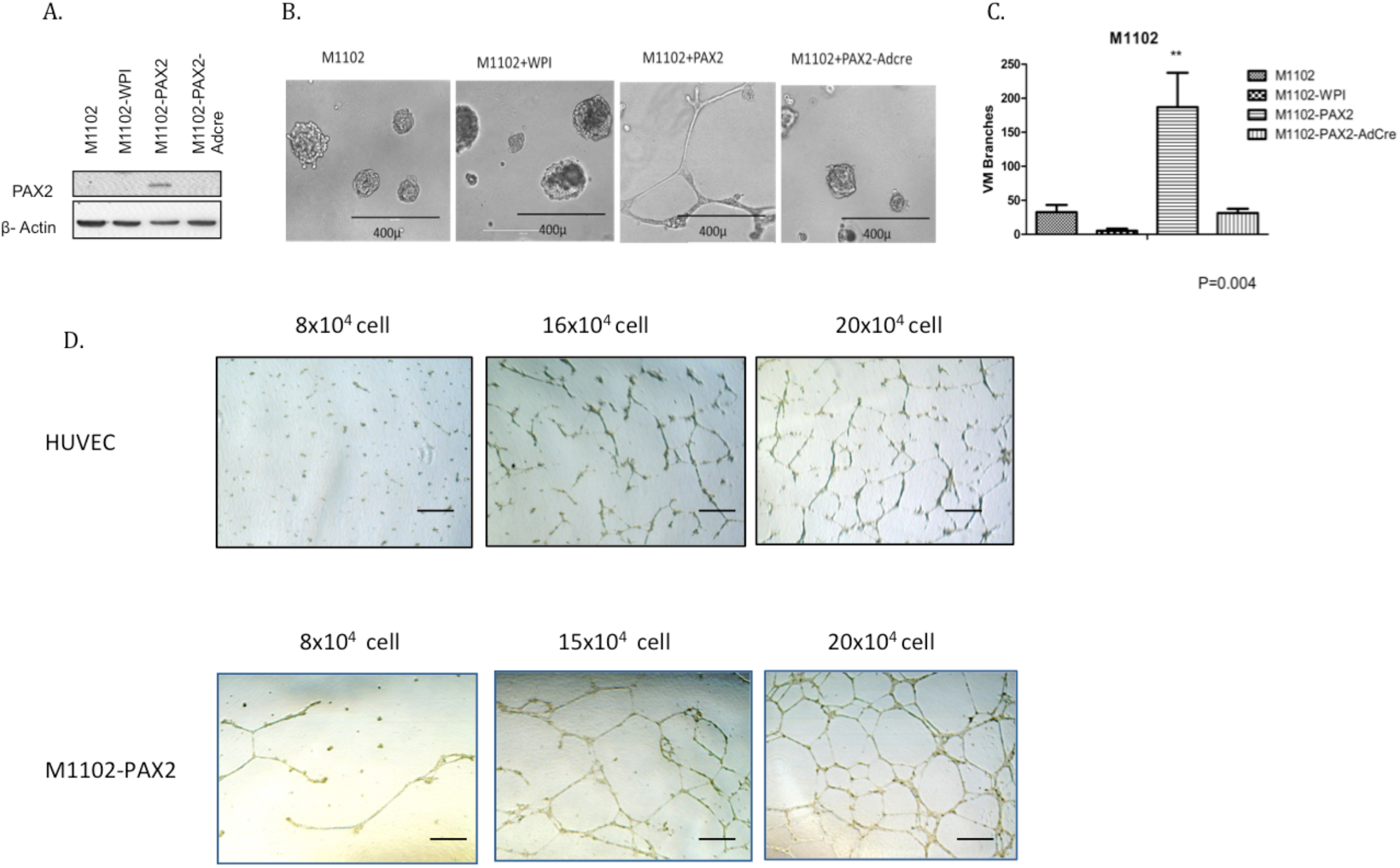
PAX2 induces the formation of vascular-like channels in 3D culture. A) Western blot shows PAX2 overexpression in MOSE cells (M1102) and knockout of PAX2 by infection with AdCre (figure previously reported (20)). B) M1102-PAX2 cells generated vascular-like channels within 48 hours, where M1102 cells without PAX2 expression developed only cell aggregations. C) Quantitative analysis of the vascular channel branches shows a significant increase in M1102-PAX2 cells compared to the control groups. Analysis was done using one-way ANOVA for three independent experiments, **p<0.01. D) The vascular channels formed by HUVEC endothelial cells and by M1102-PAX2 cells were highly dependent on cell density. The scale bar is 100μm.

### PAX2 induces endothelial independent vascular channel

To better understand if PAX2 induces epithelial cell differentiation into endothelial cells or if PAX2 induces VM, the tubular channels were stained with PAS to label the ECM formed by the M1102-PAX2 cells. The glycogen in the ECM first reacts with the periodic acid stain, which labels the tissue with dark purple color. Oxidized glycogens (aldehydes) are then detected by the Schiff stain which results in pink color in the tissue. We found that the vascular-like channels formed by M1102-PAX2 cells are PAS positive, which indicates the existence of the ECM supporting the vascular channels, whereas spheres formed by control M1102 cells are PAS negative (Fig 2A). Further, the vascular channels formed by M1102-PAX2 cells are CD31 negative, suggesting that PAX2 does not appear to induce ovarian epithelial cells to differentiate into endothelial cells, but can induce epithelial cells to form a pattern of vascular-like channels. SKOV3 cells were included as a positive control for VM since it has been previously reported that they do not express CD31 normally (13) but they are able to form VM in 3D culture (11). The SKOV3 cells readily formed vascular-like channels, and some of the cells within those channels expressed CD31 (Fig 2B). This supports previous observations of the ability of some ovarian cancer cells to acquire endothelial cell characteristics (16).

**Fig 2:**
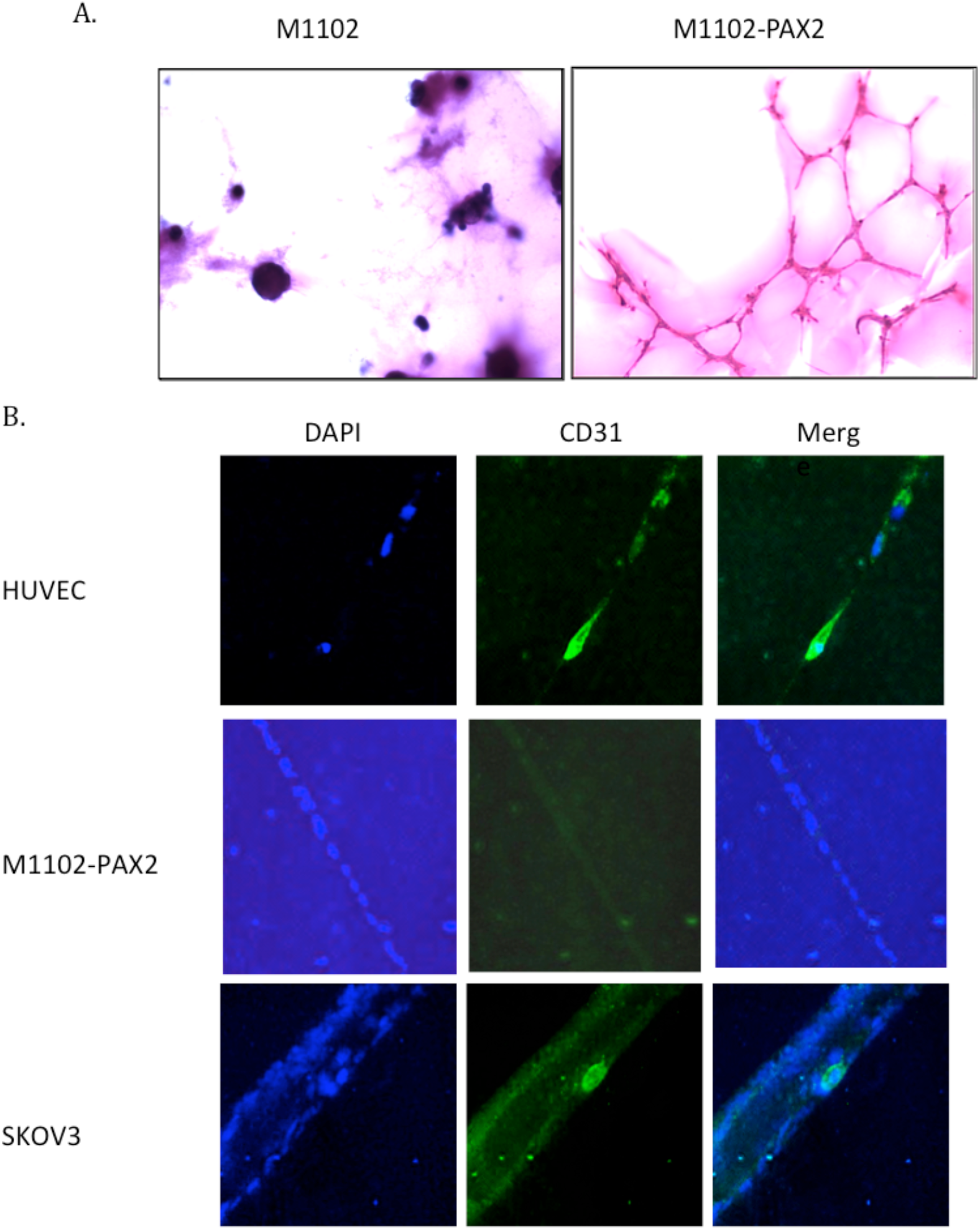
M1102-PAX2 cells formed vascular-like channels. A) PAS staining for the vascular channels formed by M1102-PAX2 cells shows pink positive staining compared to the control cells that do not express PAX2. B) Immunofluorescence staining for CD31 in the vascular channels formed by HUVEC endothelial cells, M1102–PAX2 cells and by the ovarian cancer cell line SKOV3. M1102-PAX2 vascular channels were negative for CD31 compared to HUVEC as a positive control. A few cells expressing CD31 were identified in the vascular channels formed by SKOV3 cells.

### PAX2 enhances vascular channel formation in an ovarian cancer cell line (RM)

To determine if PAX2 enhances the tumorigenesis of ovarian cancer cells by inducing VM by the cells, we analyzed the ability of PAX2 to promote VM in a mouse ovarian cancer cell line, RM. RM cells were generated by expression of activated K-RAS and c-Myc in MOSE cells and they form aggressive tumors when injected into mice (24). Overexpression of PAX2 in RM (RM-PAX2) cells resulted in even more aggressive tumors, where mice reached an average endpoint of 11 days compared to 16 days for RM cells that do not express PAX2 (25). To determine whether PAX2 might contribute to tumor progression by inducing VM, we assessed the ability of RM-PAX2 cells to form tubular channels in matrigel and compared them to the control cells (RM, RM-WPI, RM-PAX2-AdCre). The control, PAX2-non-expressing RM cells were all inefficiently able to form vascular-like channels in matrigel; however, the forced expression of PAX2 in these cells significantly enhanced the formation of vascular-like channels compared to the controls (Fig 3A & B).

**Fig 3:**
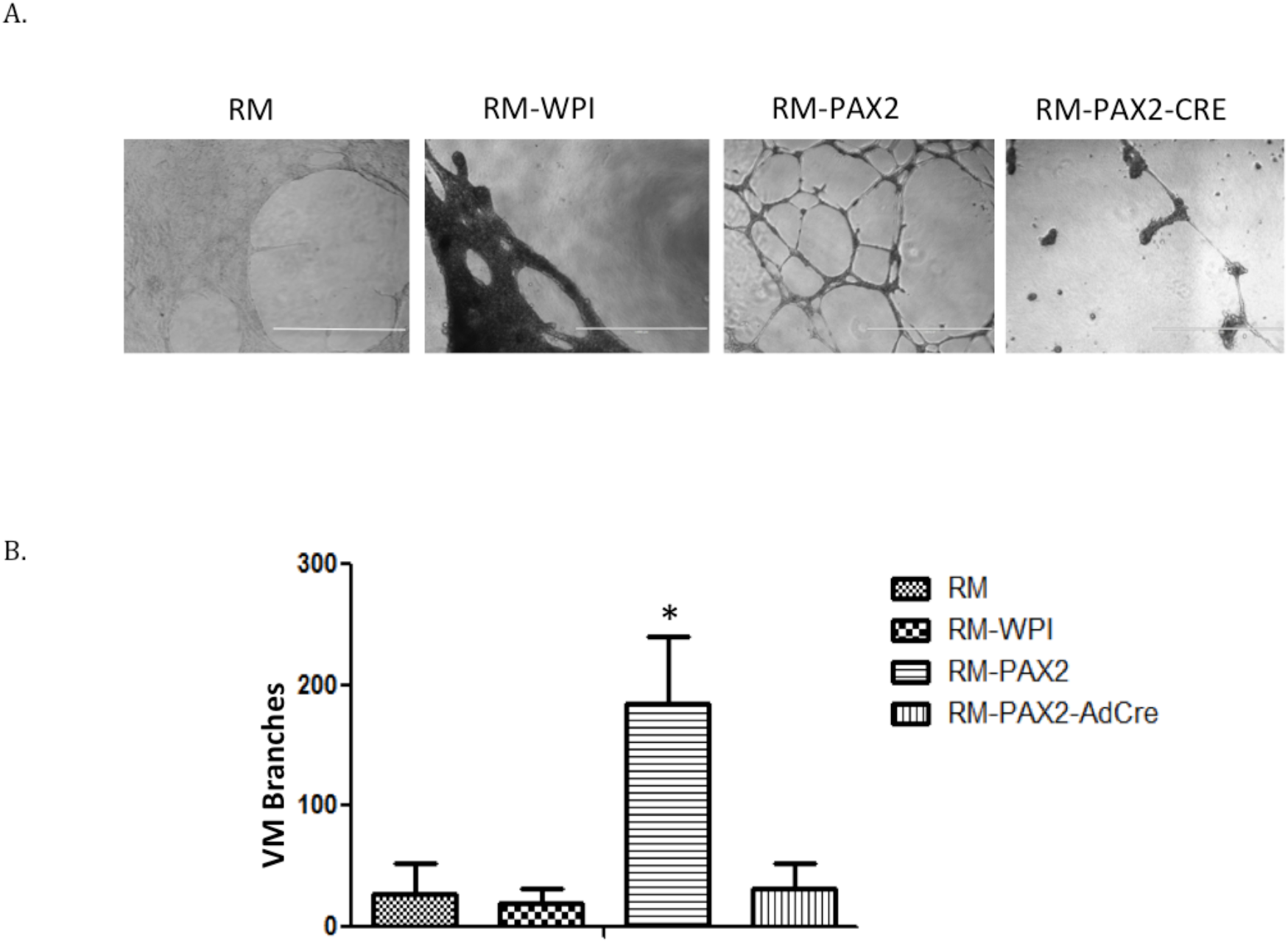
PAX2 induced formation of vascular-like channels by RM cells in 3D culture. A) RM-PAX2 cells differentiated in the matrigel to form vascular-like channels, whereas none of the control cells were able to form organized web-like tubular channels. B) Quantitative analysis of the number of vascular channel branches formed by RM-PAX2 cells shows that they are significantly more abundant than the control groups. Analysis was done using one-way ANOVA for three independent experiments, *p<0.05. The scale bar is 1000μm.

### PAX2 enhances vascular-like channel formation *in vivo* in a tumor formed by an ovarian cancer cell line

To further understand the role of PAX2 in inducing tumor progression in a model of ovarian cancer, RM cells with vector control (RM-WPI) and RM cells with forced expression of PAX2 (RM-PAX2) were injected into immune-compromised mice. Both RM-WPI and RM-PAX2 were able to develop tumors in the injected mice. The tumors were isolated, fixed and paraffin-embedded for IHC analysis. The tumor sections were stained for PAS to identify all vascular channels in the tumors, whether formed by endothelial cells or by the cancer cells. We found that RM-PAX2 tumors have plenty of small vascular channels that are PAS positive, whereas RM-WPI tumors lacked small vascular channels and PAS stain labeled primarily the large blood vessels. In addition, CD31 staining was performed on the same tumors and RM-PAX2 tumors were found to have a high density of CD31-labeled vessels compared to RM-WPI tumors, suggesting that PAX2 enhances angiogenesis in these ovarian cancers (Fig 4A & B).

**Fig 4:**
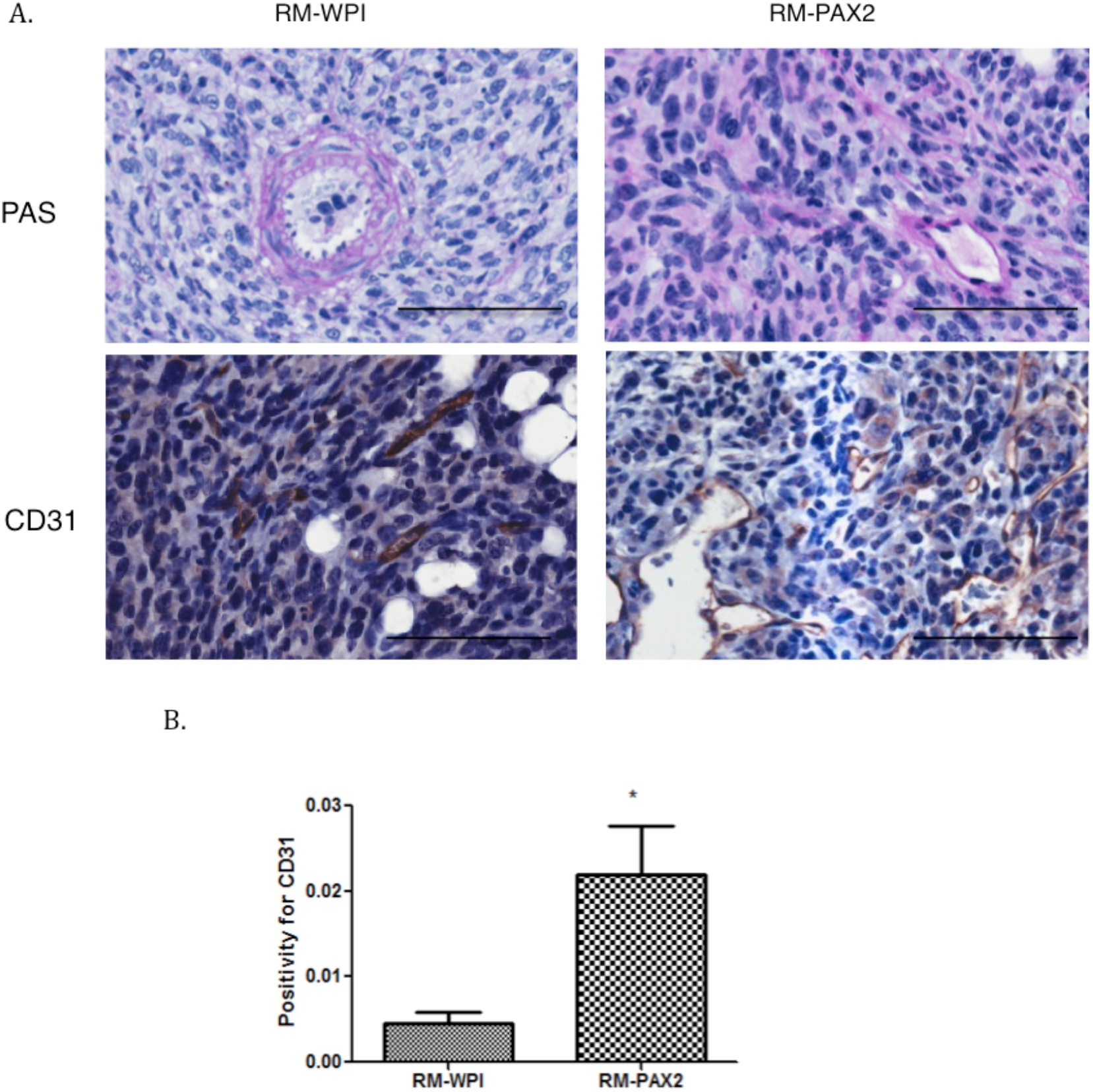
Immunohistochemistry for CD31 and PAS staining of the tumors formed by RM cells and RM cells with forced expression of PAX2. A) The upper panel of tumors derived from RM-WPI and RM-PAX2 cells shows vascular channels positive for PAS staining. The lower panel shows CD31 staining in numerous vascular channels in a RM-PAX2 tumor, indicating that PAX2 enhances the angiogenesis (yellow arrows). B) Quantification of immunohistochemical detection of CD31 shows increased CD31 positivity in RM-PAX2 compared to RM-WPI tumors. The scale bar is 100μm. Analysis was done using t-test analysis of three different fields from the tumor sections, *p<0.05.

To determine if PAX2 induces VM in ovarian tumors, we performed double staining for PAS and CD31 on the same sections. RM-PAX2 tumors had both of types of vascular channels, those formed by endothelial cells which present as PAS positive and CD31 positive, and small vascular channels that were not formed by endothelial cells, which were PAS positive and CD31 negative (Fig 5A&B). Since VM has been identified as PAS-positive/CD31-negative staining (28), these results indicate that PAX2 may enhance tumor progression by inducing VM. To confirm whether the cancer cells expressing PAX2 are the cells forming the vascular channels, we stained for PAX2 in the tumors developed by RM-WPI and RM-PAX2 cells. Tumors arising from RM-PAX2 cells had vascular channels lined with cells expressing PAX2. However, RM-WPI tumors were negative for PAX2, as expected (Fig 6A&B). These observations imply that the cancer cells expressing PAX2 can contribute to vascular channel formation in ovarian cancer.

**Fig 5:**
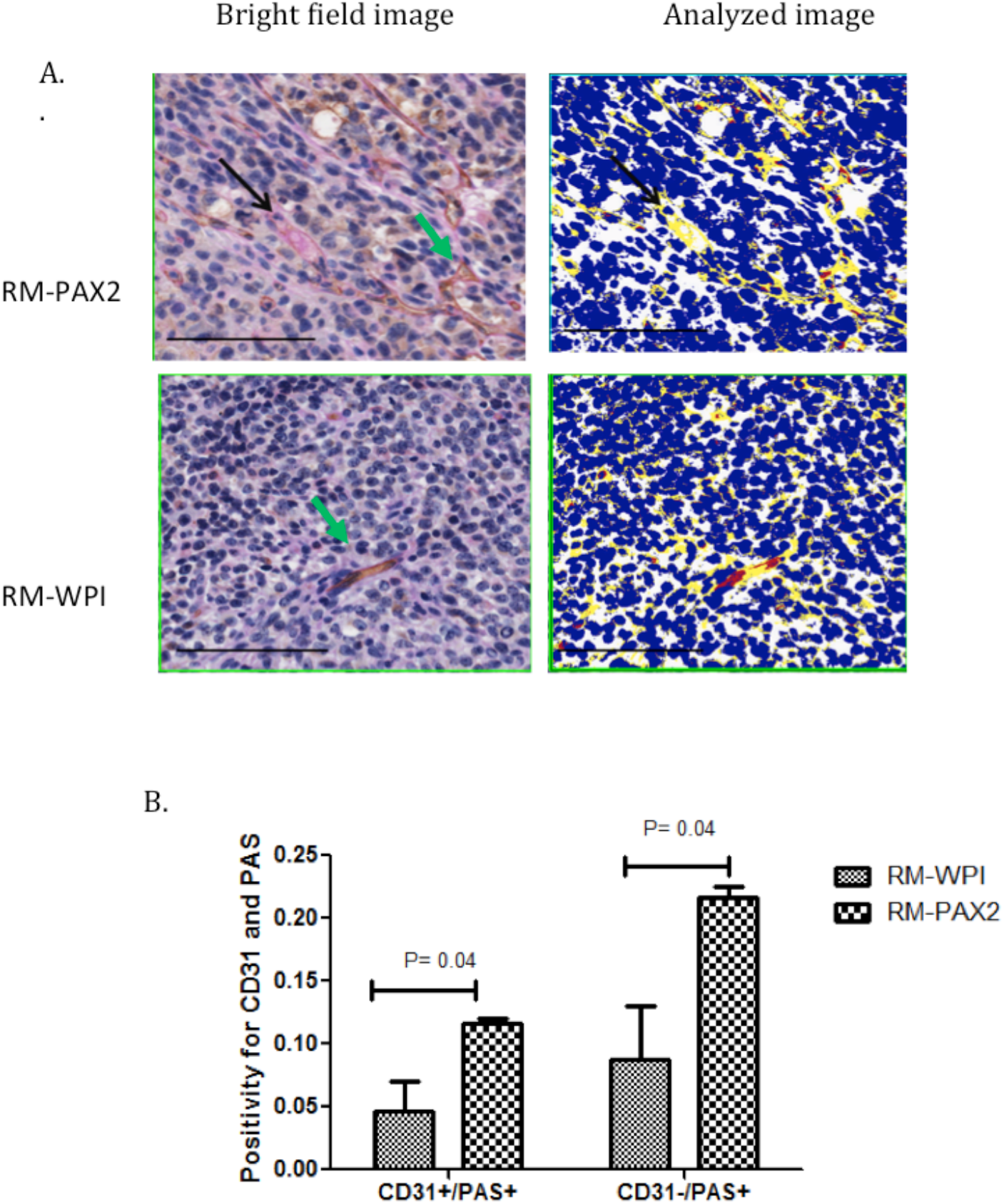
Double staining for PAS and CD31 in RM-PAX2 and RM-WPI tumors. Double staining for CD31 and PAS shows vascular channels that are PAS+/CD31+ (green arrow) in both RM-PAX2 and RM-WPI tumors, and PAS+/CD31-vascular channels (black arrow) in RM-PAX2 tumors, indicating the existence of VM in RM-PAX2 tumors. The scale bar is 100μm. B) Quantification of immunohistochemistry shows increased positivity for CD31+/PAS+ (red color) and CD31-/PAS+ (yellow color) in RM-PAX2 tumors compared to RM-WPI tumors. Image analysis was done using the Aperio Positive Pixel Count Algorithm software. Statistical analysis was done using t-test analysis of three different fields from the tumor sections.

**Fig 6:**
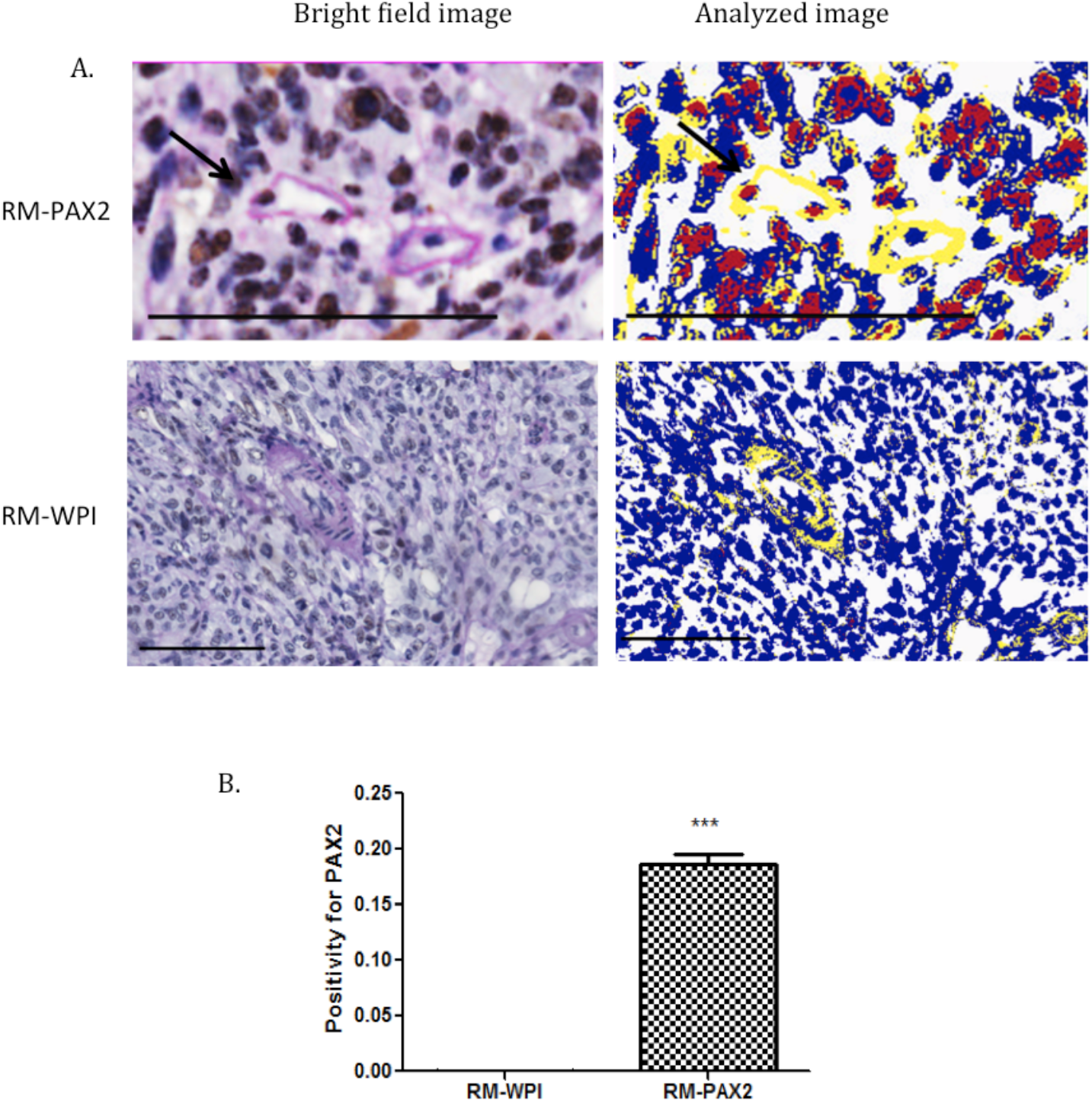
PAX2 is expressed in the cells that line the vascular channels. A) Double staining for PAS and PAX2 in RM-PAX2 and RM-WPI tumors shows, on the left, PAS-positive (pink) vascular channels with some of the cells lining the channels in RM-PAX2 tumors being positive for PAX2 (brown; arrow). The right pictures are the measurement of staining intensity; red color is PAX2 and yellow color is PAS. The scale bar is 100μm. Image Analysis was done using Aperio Positive Pixel Count Algorithm software. B) Quantification of immunohistochemical detection of PAX2 shows increased PAX2 positivity in RM-PAX2 tumors compared to RM-WPI tumors. The quantification of PAX2 expression was determined based on positive pixel counts relative to the tissue area. Analysis was done using t-test analysis for three different fields from the tumor sections. ***p<0.001.

## Discussion

The role of PAX2 in ovarian cancer is poorly understood, but the results of previous studies suggested that expression of PAX2 alone is not able to induce OSE cell transformation into cancer (25). However, the ability of PAX2 to promote VM suggests that tumors that express PAX2 might be more aggressive through their development of VM in tumors, thereby augmenting tumor blood supply without angiogenesis.

Normal OSE cells did not express PAX2 (25), whereas 61% of ovarian cancer cell lines express PAX2 (17), suggesting a potential oncogenic role for PAX2 in cancers derived from the ovarian or fallopian tube epithelium. The oncogenic role of PAX2 in ovarian cancer has recently been investigated in terms of its ability to promote cell proliferation, migration and tumor formation (25). In our attempts to confirm the ability of PAX2 to promote epithelial differentiation of OSE cells, we found that PAX2 expression in M1102 OSE cells induced the formation of tubular-like structures in matrigel, whereas the control groups did not form these structures. The ability to form these tubular or vessel-like structures has been previously associated with aggressive tumor cell lines such as melanoma (3) and the phenomena is known as vasculogenic mimicry (VM). While the formation of vascular networks by normal endothelial cells is a standard measure of their organizational potential (29), no other normal cells have been reported to have this capability. Aggressive cancer cell lines, however, can organize in a pattern to form *de novo* these webs of vessels (3). The tubular networks formed by M1102-PAX2 in matrigel were morphologically similar to the vessels formed by HUVEC cells and to the VM networks formed by SKOV3 ovarian cancer cells.

The M1102-PAX2 tubular webs were supported by remodeling ECM which was positive for PAS staining and negative for CD31. However, SKOV3 VM had a few cells expressing CD31, which is consistent with previous reports suggesting that cancer cells may differentiate into endothelial cells *de novo* to induce the angiogenesis (16). Thus, we hypothesized that PAX2 induces VM *in vitro* in normal MOSE cells. VM induction in M1102 by PAX2 is not a result of stem cell differentiation since PAX2 decreases stemness in MOSE cells, as we previously reported (20). Unfortunately, we were not able to investigate the vessel formation *in vivo* with normal MOSE-PAX2, since these cells are not able to form tumors when they have been injected into mice (25).

To further investigate the possible role of PAX2 in VM during tumor progression, we used a mouse model of ovarian cancer cells (RM) created by transformation of MOSE cells by *K-Ras* and *c-Myc* mutations. The forced expression of PAX2 in RM cells significantly increased the tubular formation *in vitro*. RM cells have been shown to be tumorigenic in mice and the forced expression of PAX2 induced more rapid tumor progression (25). To understand if PAX2 expression enhances vasculogenesis by the cancer cells *in vivo*, the tumors formed by RM and RM-PAX2 cells were analyzed for PAX2, CD31 expression and for PAS staining. RM-PAX2 increases angiogenesis as determined by staining the tumor with CD31. However, in the same tumor, some of the neo-vasculatures were negative for CD31 and positive for PAS staining which indicates that endothelial cells did not form these new vessels. Interestingly, some of the cells lining those vessels expressed PAX2 in the nucleus, which indicates that cancer cells that have high expression of PAX2 can contribute to the formation of neo-vasculature in the tumor. The molecular mechanism by which PAX2 can promote VM is an interesting question to be addressed in future studies. Since we know that PAX2 induces COX2 in RM cells (25) and COX2 has been reported to induce VM in breast cancer (10), we anticipate that PAX2 induced VM may be mediated by COX2 expression.

The potential role for PAX2 in VM is novel, but PAX2 has been reported to induce tubular branching in the kidney and to induce the elongation of the inner ear precursors (30) and it may play a similar role in fallopian tube morphogenesis (25). In established tumors, PAX2 may enhance tumor progression and metastasis by vasculogenic mimicry, which enables increased blood supply for the tumor. However, the role of PAX2 in inducing cell branching or tubular formation and the protein expression that induces VM in the ovarian cancer warrants further investigation.

## Acknowledgments

The authors thank Olga Collins for her excellent technical support.

## References

1. Piao Y, Liang J, Holmes L, Henry V, Sulman E, de Groot JF. Acquired resistance to anti-VEGF therapy in glioblastoma is associated with a mesenchymal transition. Clin Cancer Res. 2013 Aug;19(16):4392–403.

2. Helfrich I, Scheffrahn I, Bartling S, Weis J, von Felbert V, Middleton M, et al. Resistance to antiangiogenic therapy is directed by vascular phenotype, vessel stabilization, and maturation in malignant melanoma. J Exp Med. 2010 Mar;207(3):491–503.

3. Maniotis AJ, Folberg R, Hess A, Seftor EA, Gardner LM, Pe’er J, et al. Vascular channel formation by human melanoma cells in vivo and in vitro: vasculogenic mimicry. Am J Pathol. 1999 Sep;155(3):739–52.

4. Racordon D, Valdivia A, Mingo G, Erices R, Aravena R, Santoro F, et al. Structural and functional identification of vasculogenic mimicry in vitro. Sci Rep. 2017;7(1).

5. Valdivia A, Mingo G, Aldana V, Pinto MP, Ramirez M, Retamal C, et al. Fact or Fiction, It Is Time for a Verdict on Vasculogenic Mimicry? Front Oncol. 2019;9.

6. Itzhaki O, Greenberg E, Shalmon B, Kubi A, Treves AJ, Shapira-Frommer R, et al. Nicotinamide inhibits vasculogenic mimicry, an alternative vascularization pathway observed in highly aggressive melanoma. PLoS One. 2013;8(2):e57160.

7. Dong X, Sun B, Zhao X, Liu Z, Gu Q, Zhang D, et al. Expression of relative-protein of hypoxia-inducible factor-1alpha in vasculogenesis of mouse embryo. J Biol Res (Thessalonike, Greece). 2014 Dec;21(1):4.

8. Luo F, Yang K, Liu R-L, Meng C, Dang R-F, Xu Y. Formation of vasculogenic mimicry in bone metastasis of prostate cancer: correlation with cell apoptosis and senescence regulation pathways. Pathol Res Pract. 2014 May;210(5):291–5.

9. Qiao L, Liang N, Zhang J, Xie J, Liu F, Xu D, et al. Advanced research on vasculogenic mimicry in cancer. J Cell Mol Med. 2015 Feb;19(2):315–26.

10. Basu GD, Liang WS, Stephan DA, Wegener LT, Conley CR, Pockaj BA, et al. A novel role for cyclooxygenase-2 in regulating vascular channel formation by human breast cancer cells. Breast Cancer Res. 2006;8(6):R69.

11. Millimaggi D, Mari M, D’Ascenzo S, Giusti I, Pavan A, Dolo V. Vasculogenic mimicry of human ovarian cancer cells: role of CD147. Int J Oncol. 2009 Dec;35(6):1423–8.

12. Sun J, Hemler ME. Regulation of MMP-1 and MMP-2 production through CD147/extracellular matrix metalloproteinase inducer interactions. Cancer Res. 2001 Mar;61(5):2276–81.

13. Sood AK, Seftor EA, Fletcher MS, Gardner LM, Heidger PM, Buller RE, et al. Molecular determinants of ovarian cancer plasticity. Am J Pathol. 2001 Apr;158(4):1279–88.

14. Ayala-Domínguez L, Olmedo-Nieva L, Muñoz-Bello JO, Contreras-Paredes A, Manzo-Merino J, Martínez-Ramírez I, et al. Mechanisms of Vasculogenic Mimicry in Ovarian Cancer. Front Oncol. 2019;9.

15. Tang HS, Feng YJ, Yao LQ. Angiogenesis, vasculogenesis, and vasculogenic mimicry in ovarian cancer. Vol. 19, International Journal of Gynecological Cancer. 2009. p. 605–10.

16. Su M, Feng Y-J, Yao L-Q, Cheng M-J, Xu C-J, Huang Y, et al. Plasticity of ovarian cancer cell SKOV3ip and vasculogenic mimicry in vivo. Int J Gynecol Cancer. 2008;18(3):476–86.

17. Song H, Kwan S-Y, Izaguirre DI, Zu Z, Tsang YT, Tung CS, et al. PAX2 Expression in Ovarian Cancer. Int J Mol Sci. 2013;14(3):6090–105.

18. Eccles MR, He S, Legge M, Kumar R, Fox J, Zhou C, et al. PAX genes in development and disease: the role of PAX2 in urogenital tract development. Int J Dev Biol. 2002;46(4):535–44.

19. Terzić J, Muller C, Gajović S, Saraga-Babić M. Expression of PAX2 gene during human development. Int J Dev Biol. 1998;42(5):701–7.

20. Alwosaibai K, Abedini A, Al-Hujaily EM, Tang Y, Garson K, Collins O, et al. PAX2 maintains the differentiation of mouse oviductal epithelium and inhibits the transition to a stem cell-like state. Oncotarget. 2017;8(44):76881–97.

21. Narlis M, Grote D, Gaitan Y, Boualia SK, Bouchard M. Pax2 and pax8 regulate branching morphogenesis and nephron differentiation in the developing kidney. J Am Soc Nephrol. 2007 Apr;18(4):1121–9.

22. Mansouri a, Hallonet M, Gruss P. Pax genes and their roles in cell differentiation and development. Curr Opin Cell Biol. 1996;8(6):851–7.

23. McCloskey CW, Goldberg RL, Carter LE, Gamwell LF, Al-Hujaily EM, Collins O, et al. A new spontaneously transformed syngeneic model of high-grade serous ovarian cancer with a tumor-initiating cell population. Front Oncol. 2014;4:53.

24. Yao D-S, Li L, Garson K, Vanderhyden BC. [The mouse ovarian surface epithelium cells (MOSE) transformation induced by c-myc/K-ras in]. Zhonghua Zhong Liu Za Zhi. 2006 Dec;28(12):881–5.

25. Al-Hujaily EM, Tang Y, Yao D-S, Carmona E, Garson K, Vanderhyden BC. Divergent roles of PAX2 in the etiology and progression of ovarian cancer. Cancer Prev Res (Phila). 2015 Sep;

26. Kutner RH, Zhang X-Y, Reiser J. Production, concentration and titration of pseudotyped HIV-1-based lentiviral vectors. Nat Protoc. 2009;4(4):495–505.

27. Debnath J, Muthuswamy SK, Brugge JS. Morphogenesis and oncogenesis of MCF-10A mammary epithelial acini grown in three-dimensional basement membrane cultures. Methods. 2003 Jul;30(3):256–68.

28. Folberg R, Hendrix MJC, affiliation> AJM. Vasculogenic mimicry and tumor angiogenesis. Am J Pathol [Internet]. 2000;156(2):361–81. Available from: http://dx.doi.org/10.1016/S0002-9440(10)64739-6

29. Ponce ML. Tube formation: an in vitro matrigel angiogenesis assay. Methods Mol Biol. 2009;467:183–8.

30. Christophorou N a D, Mende M, Lleras-Forero L, Grocott T, Streit A. Pax2 coordinates epithelial morphogenesis and cell fate in the inner ear. Dev Biol [Internet]. 2010;345(2):180–90. Available from: http://dx.doi.org/10.1016/j.ydbio.2010.07.007

